# Multi-GBS: A massively multiplexed GBS-based protocol optimized for large, repetitive conifer genomes

**DOI:** 10.1101/2022.01.10.475685

**Authors:** Miguel Vallebueno-Estrada, Sonja Steindl, Vasilina Akulova, Julia Riefler, Lucyna Slusarz, Alexis Arizpe, Kelly Swarts

## Abstract

Reduced representation library approaches are still a valuable tool for breeding and population and ecological genomics, even with impressive increases in sequencing capacity in recent years. Unfortunately, current approaches only allow for multiplexing up to 384 samples. To take advantage of increased sequencing capacity, we present Multi-GBS, a massively multiplexable extension to Genotyping-by-Sequencing that is also optimized for large conifer genomes. In Norway Spruce, a highly repetitive 20Gbp diploid genome with high population genetic variation, we call over a million variants in 32 genotypes from three populations, two natural forest in the Alps and Bohemian Alps, and a managed population from southeastern Austria using the existing TASSEL GBSv2 pipeline. Metric MDS analysis of replicated genotypes shows that technical bias in resulting genotype calling is minimal and that populations cluster in biologically meaningful ways.

## Introduction

While whole-genome sequencing of diverse populations is becoming increasingly feasible, there is still a need for reduced representation approaches for species with large genomes and for very large-scale population sequencing in breeding and population genomics. Reduced representation approaches maximize read overlap across populations while still mapping to a small fraction of the genome, saving costs. The most popular reduced representation approaches include capture methods (Ali et al., 2016; Andermann et al., 2019; Bernhardsson et al., 2020; Carpenter et al., 2013), where “bait” sequences are hybridized to target DNA molecules and isolated for sequencing, and restriction digest approaches (Elshire et al., 2011; Etter et al., 2011; Wallace & Mitchell, 2017) that target restriction cut site sequences to anchor reads.

Both of these strategies are effective for generating comparable variants across populations, but restriction digest approaches are appealing for high diversity population genomics because they are unbiased with respect to the ascertainment biases of the population that generated the bait sequences (Heslot et al., 2013). Additionally, they are lower cost than synthesized capture probes and will not be depleted, which can be a problem in probes generated from biological samples. Unfortunately, the published reduced representation protocols are unsuitable for some species such as conifer systems because they fail to produce replicable libraries without heavy filtering (Xiao-ru Wang, personal communication, 2019). More generally, it is currently not possible to multiplex more than 384 samples, a barrier to cost-effectively using new short-read technologies. We present here a highly replicable, massively-multiplexable, robust GBS-based protocol tested on Norway Spruce (*Picea abies*), a highly diverse conifer with a repetitive, 20 Gbp genome.

## Materials and Methods

### Library construction design

Published GBS library protocols (Elshire et al., 2011) (Wallace & Mitchell, 2017) served as baseline for the protocol. The full protocol is available as supporting information. Deviations are as follows:

1. As in the original protocol, we use the methyl sensitive endonuclease ApeKI (NEB R0643L) in order to minimize the repetitive fraction sampled, while still targeting a large fraction of the accessible (functional) regions of the genome. To adjust for this, starting DNA was increased from 100 ng to 150 ng (Figure 1 step 1 and 2).
2. Because we found restriction digestion to be sometimes incomplete at 2 hours, we increased the digestion to overnight (14 hours) (Figure 1 step 2). NEB buffer 3.1 at the time of testing was unstable. Always check that the digested product follows the expected fragment distribution profile.
3. Ligation was increased to 1.5 hours to ensure complete ligation of fragments (Figure 1 step 3).
4. Because the fragment distribution of spruce was shifted to larger digested fragments relative to maize, we use a Phusion high-fidelity polymerase rather than the Taq that is used in the traditional protocol. This necessitates changing PCR conditions. We test here a 15 and 30 second extension time, leading to an expected exponential increase of fragments up to 1000 and 2000 bp respectively. Cycle number was reduced from 18 to 13 to retain library complexity (Figure 1 step 5).
5. Magnetic bead cleanups were used instead of column cleanups to enable plate-based robotics. (Figure 1 steps 4 and 6).
6. The volume of the restriction digest and ligation reaction is decreased by half to save costs. Total reagent costs/sample are less than one euro.
7. We modify the illumina adapters as follows to add TruSeq UDI p5 and p7 barcoded adapters. We use the internal barcodes to identify wells of each plate, but the addition of the TruSeq adapters allows us to uniquely identify up to 96 plates multiplexed in a sequencing lane. We created two new adaptors: one common adaptor P7 includes the reverse complement sequence of the sequencing primer 2 of the TrueSeq UDI and added a sticky end on the 3’ complementary to the restriction pattern of Apk1 (Figure 1 step 3). The adaptor P5 is the reverse complement sequence of the sequencing primer 2 of the TruSeq UDI, where we included a GBS tag and a sticky end on the 5’ complementary to the restriction pattern of Apk1. Sequencing Truseq indexes and flow cell complementary sequences are added during PCR by two primers complementary to the Truseq UDI adapter sequences (Figure 1 step 5.)

**Figure 1.**
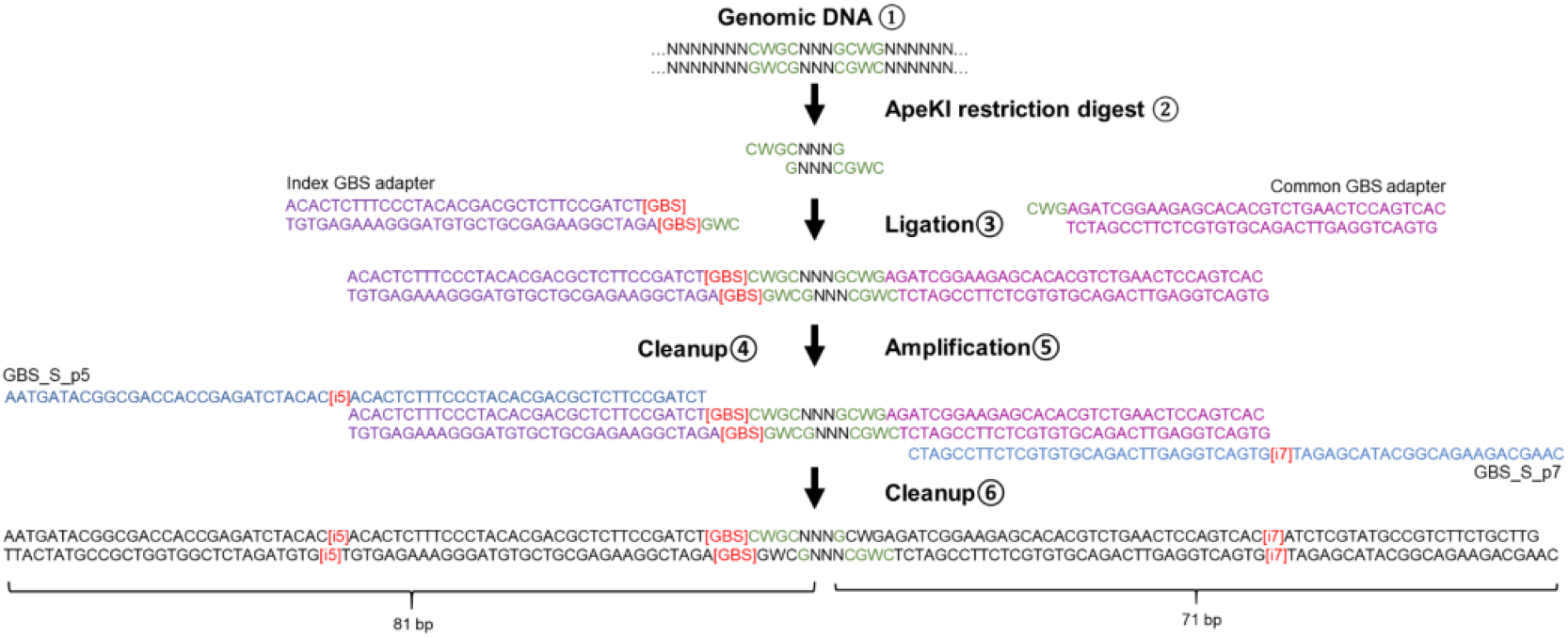
Multi-GBS library construction design. Anchor ApeKI restriction sites are colored in green, Common GBS double stranded adapter is colored in magenta, index GBS double stranded adapter is colored in purple. Amplification primers GBS_S_p5 and GBS_Sp7 ar e depicted in blue. The legend [GBS] in red represents an index sequence for the GBS pipeline. The legends [i5] and [i7] in red represent the P5 and P7 dual sequencing indexes respectively.

**Figure 2.**
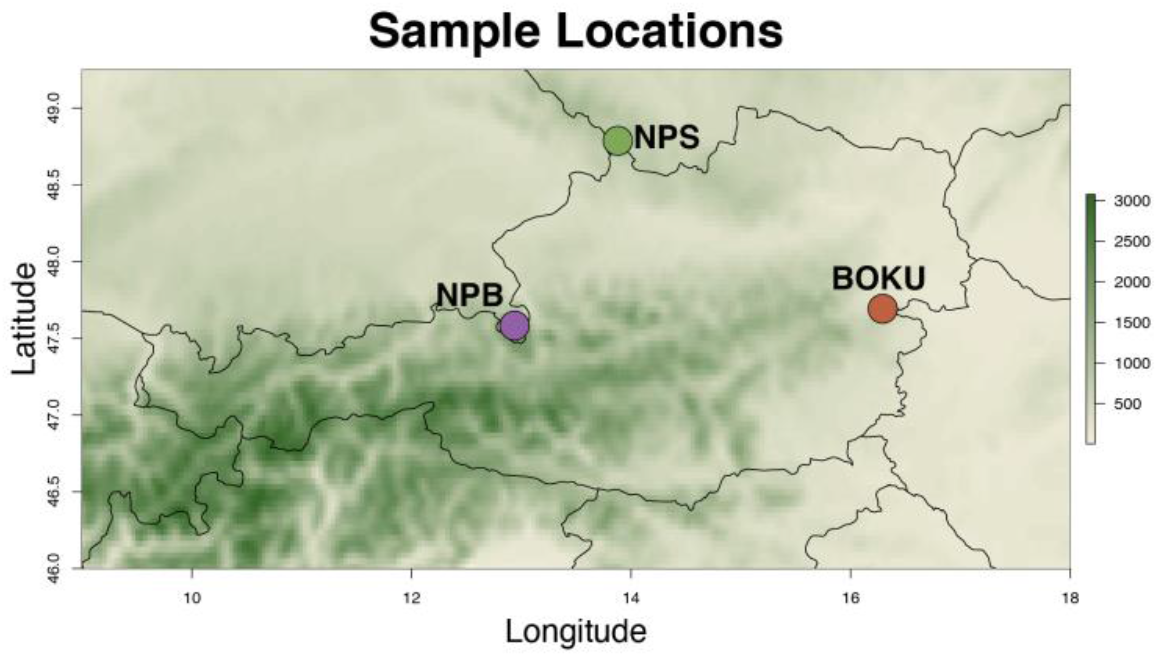
Sampling locations for testing

**Figure 3.**
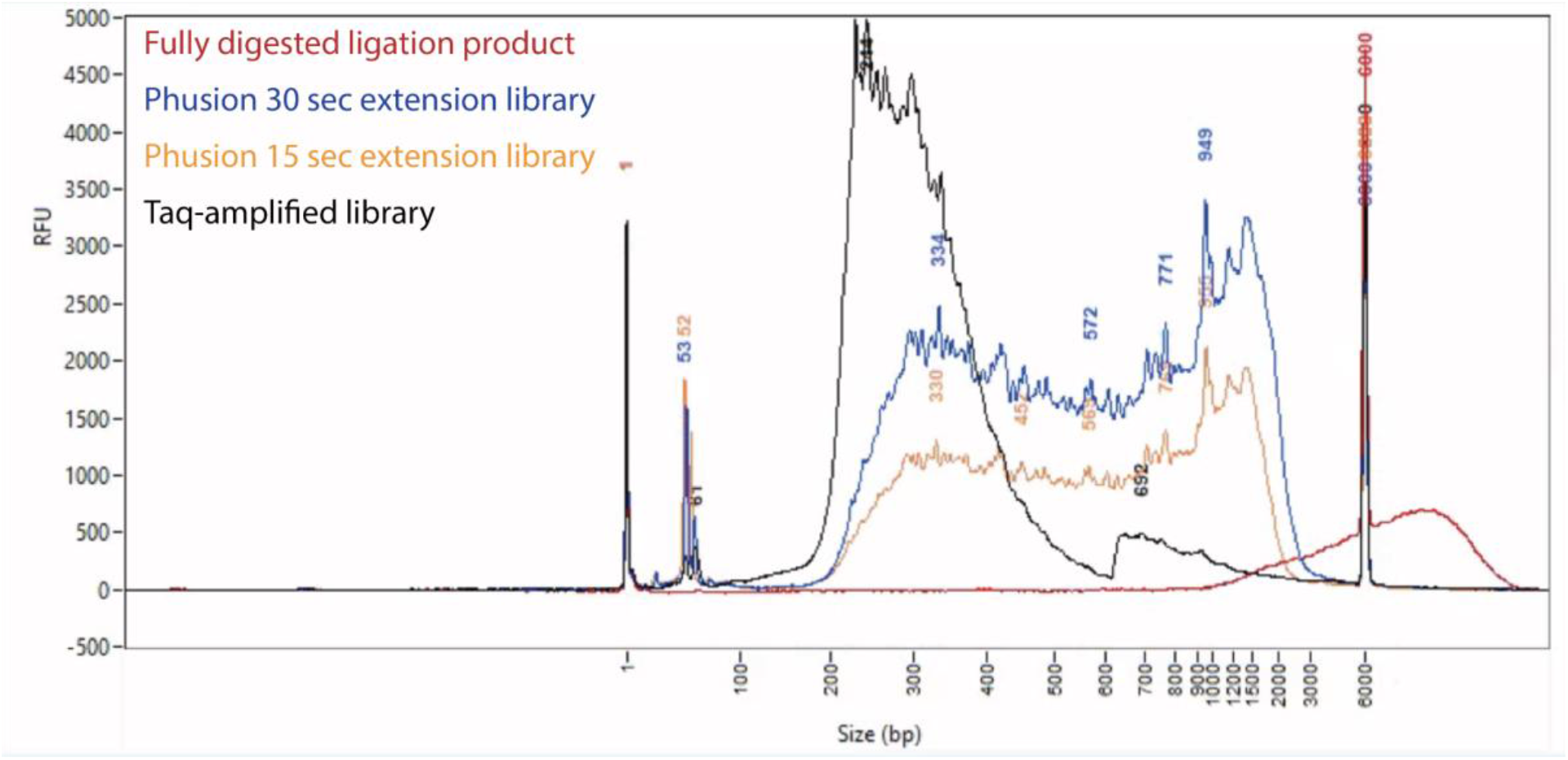
Restriction fragment profiles for restriction digest pattern in P. abies, and resulting libraries amplified with Taq and Phusion polymerases

### Protocol testing for technical replicability, variant profiling and population discrimination

#### Tissue sampling

Cambial tissue samples were collected under permit from Berchtesgaden National Park (BNP), Germany, Šumava National Park (SNP), Czechia and the University of Natural Resources and Life Sciences teaching forest in Burgenland, Austria (BOKU) over the summer of 2021. 10-11 samples were collected in each location, which represents natural forest populations in the Alps (NPB),

Bohemian Alps (NPS) and an isolated, managed forest in the tail of the Alps (BOKU). Tissue was collected with a 5 mm increment core borer and trimmed with clippers into a qiagen extraction plate with 600 ul of 100% ethanol. In the lab, ethanol was removed and tissue was dried in a lyophilizer and stored at -20C.

#### DNA Extraction

DNA was extracted at the Gregor Mendel Institute following a cambial CTAB (Doyle & Doyle, 1987) extraction protocol modified from the Aitken group (Hanlon, 2017). This protocol yields high quality DNA with a median fragment length of 125 kbp.

#### Library Preparation to test digestion

150 ng of each sample randomized across two plates previously prepared with 3.6 ng of unique internally barcoded GBS adapters, replicated twice per plate, and dried down in a SpeedVac at 30°C. The plate digested for 14 hours was amplified twice with unique TruSeq adapter pairs, once with a 15 second extension time and the other with a 30 second extension time to test. The Phusion polymerase and buffer used were produced in-house at the Vienna BioCenter Molecular Biology Services.

#### Sequencing

Both plates of libraries were sequenced together in a single partial lane of an Illumina NovaSeqS4 on paired-end 150 bp mode at the ViennaBioCenter Sequencing Facility. This allows us to test the benefit of longer read sequencing. Variants were called using only the forward strand and the incomplete digest library was very low concentration.

#### Variant calling

Variant calling was done according to the recommended TASSEL GBSv2 pipeline (Bradbury et al., 2007; *Tassel5GBSv2Pipeline*, 2021) using default values with the following exceptions:

- GBSSeqToTagDBPlugin -c 2 -kmerLength 64<or>120 -mnQS 20 -mxKmerNum 300000000
- TagExportToFastqPlugin -c 1 -endPlugin -runfork1 & > “$dir/TagExportToFastqPlugin.log”; SAMToGBSdbPlugin -minMAPQ 20 -mapper bwaMem
- DiscoverySNPCallerPluginV2 -mnLCov 0.1 -mnMAF 0.01
- ProductionSNPCallerPluginV2 -kmerLength 64<or>120 -mnQS 20

Mapping was performed using bwa mem against versions of the *Picea abies* reference genomes downloaded from ConGenIE (Nystedt et al., 2013): “Pabies1.0-all-cds”, “Pabies1.0-genome” and an additional reference we generated by subsetting the “Pabies1.0-genome” reference for only the contigs with cds regions, as defined in the contig names of “Pabies1.0-all-cds”. This reference is called “Pabies1.0-genome-with-cds”. Contigs from each of these three references were concatenated together with 100 “N”s dividing each region to facilitate SNP calling and resulting variants were reassigned to their original contig based on an associated positional lookup file. The perl script to concatenate the contigs and generate the lookup is available as a supporting resource.

Six keysets controlling the generation of 36 unique variant call datasets were generated. Each independent library is unique in the “All” dataset. In “15s” and “30s” only the 15 second and 30 second, respectively, extension samples were used. Each of these sets were called merging replicate genotypes as “MergeAll”, “Merge15s” and “Merge30s” respectively. Each of these six sets were mapped against the three reference genomes using the kmer settings “60bp” and “120bp”.

#### Metric MDS and variant analyses for genome coverage and ensuring technical replicability

IBS (Hamming) distance matrices were generated in TASSEL (Bradbury et al., 2007) after first randomly downweighting heterozygotes with a probability of 0.5. Only taxa with greater than 20% coverage were included to ensure sufficient pairwise comparisons for accurate distance estimation. Metric MDS was performed using cmdscale in R based on the distance matrices. Downweighting heterozygotes accounts for SNP undercalling due to variable coverage but also has the effect of increasing genetic distance between replicate samples as a function of their true heterozygosity. Results can be interpreted similarly to principal components analysis.

## Results

### Restriction enzyme digest and library profiles in *P. abies*

The digested fragment distribution of *P. abies* is overwhelmingly greater than 1kbp, making Taq, a slow, error-prone polymerase, less effective for amplifying diverse fragments than the fast, high-fidelity Phusion. The average size of the 30s library is about 100bp larger than the 15s library, suggesting that it does not sample substantially more fragments and should be expected to have a higher proportion of linearly-amplified fragments.

### Library characteristics

Phusion libraries yield less than Taq with about 5ng/ul on average for Phusion vs arround 20ng/ul for Taq, which is expected due to the reduction in cycle number from 18 to 13. The 15 sec extension plate with 762,282,481 reads and the 30 sec plate with 678,301,151 reads. Mean “good barcoded read” (*Tassel5GBSv2Pipeline*, 2021) - reads with a residual cut site and no Ns within the length of the kmer (either longer than the kmer length or runs into the illumina adapter) - proportion is 0.48 for the 64-mer and 0.38 for the 120-mer and mean “low quality read” - reads where any of the base pair qualities fall below the designated value within the barcode plus kmer region - is 0.13 for 64-mer and 0.24 for 120-mer. The low quality reads can be minimized by decreasing the “-mnQS” flag, especially when using illumina technologies with a uniform error profile. Proportion good barcoded reads was lower for 120 kmer length than for 64 at both 15 seconds (0.49 vs 0.37) and 30 second (0.46 vs 0.36) and 30 second extension was lower than 15 seconds, suggesting that there is a significant population of incompletely amplified fragments in the library. This is likely due to the variability of the fragment length and the long extension time, where short fragments may finish early and start another. These fragments could be effectively reduced in the final library by performing a size selection to adapter length (152bp)+read length. The effect is species dependent, but based on the fragment distribution in *P. abies*, very few reads would be lost with a final size selection for molecules greater than 300 bp.

### Variant characterization

Up to 1.6M variants were called across the genomic contigs using only 32 unique but technically replicated genotypes in 120-mer mode. Even using unreplicated genotypes, up to 1.2M variants were called. Mean depth of coverage for individual libraries (All,15s,30s) called on an average of 4.5M 120-mers per sample was 11.5 across the genome and 18 in the CDS and 16 for contigs with CDS. Based on an average of 5.8M 64-mers from the same reads, mean depth of coverage was 16 across the genome and 29 in the CDS and 26 for contigs with CDS.

**Table 1.** Variant information for datasets. Reference genome aligned to represented as “genomic contigs with CDS regions/contigs for CDS regions/genomic contigs”

| 120-mer |  |  |  |  |  |  |
| --- | --- | --- | --- | --- | --- | --- |
| **All** | 0.74/0.53/0.04 | 20/8/2 | 37.19/16.59/3.57 | 0.67/0.61/0.58 | 0.14/0.17/0.08 | 1,637,415/621,531/1,622,796 |
| **15s** | 0.7/0.48/0.04 | 16/6/2 | 29.25/14.28/3.13 | 0.72/0.66/0.61 | 0.17/0.2/0.11 | 1,224,874/492,065/1,153,365 |
| **30s** | 0.69/0.47/0.03 | 14/6/2 | 27.09/13.71/3.06 | 0.65/0.7/0.59 | 0.18/0.2/0.1 | 1,105,125/455,988/1,025,650 |
| **MergeAll** | 0.75/0.52/0.04 | 19/7/2 | 35.43/15.57/3.28 | 0.8/0.83/0.78 | 0.23/0.26/0.15 | 1,476,036/1,573,034/459,552 |
| **Merge15** | 0.7/0.48/0.03 | 15/6/2 | 27.81/13.58/2.98 | 0.78/0.82/0.74 | 0.23/0.26/0.15 | 1,156,751/459,552/1,048,482 |
| **Merge30** | 0.68/0.46/0.03 | 13/6/2 | 25.81/13.07/2.91 | 0.77/0.81/0.73 | 0.23/0.26/0.15 | 1,045,184/426,334/937,392 |
| 64-mer |  |  |  |  |  |  |
| **All** | 0.71/0.46/0.03 | 12/5/2 | 21.66/9.38/2.76 | 0.64/0.71/0.61 | 0.12/0.15/0.07 | 912,797/303,832/890,837 |
| **15s** | 0.67/0.41/0.03 | 9/4/2 | 17.11/8.24/2.43 | 0.68/0.75/0.64 | 0.15/0.18/0.09 | 680,880/242,837/636,796 |
| **30s** | 0.65/0.4/0.02 | 9/4/1 | 15.78/7.93/2.37 | 0.67/0.74/0.63 | 0.15/0.18/0.09 | 225,013/612,453/566,515 |
| **MergeAll** | 0.71/0.44/0.03 | 11/4/2 | 20.55/8.76/2.58 | 0.81/0.86/0.81 | 0.2/0.24/0.14 | 869,120/277,030/795,071 |
| **Merge15** | 0.66/0.4/0.02 | 9/4/1 | 16.17/7.83/2.33 | 0.8/0.85/0.77 | 0.2/0.23/0.14 | 638,032/224,087/574,516 |
| **Merge30** | 0.64/0.39/0.02 | 8/4/1 | 15/7.53/2.27 | 0.79/0.84/0.76 | 0.21/0.24/0.14 | 575,620/207,927/513,215 |
|  | Contig coverage | Median SNPs/covered contig | Mean SNPs/covered contig | Median taxa coverage | Median heterozygosity | N variants |

For unique libraries, variants were called on 74% of contigs with CDS regions, with a median of 20 SNPs per contig and 53% of the CDS regions themselves had variants called with a median of 8 SNPs/region. As is expected with a methyl sensitive enzyme, only 4% of genomic contigs contained variants with a median of 2 SNPs/contig. Median taxa coverage for unique libraries genome-wide was 67% with 14% heterozygosity, increasing to 80% with 23% heterozygosity when genotypes were merged. The difference is lower when only the 15s or 30s libraries alone were used, as might be expected given that they sample slightly different genomic space. Heterozygosity improves somewhat in the merged samples, suggesting that even with the high depth of coverage, not all loci are saturated.

### Replicability and ability to discriminate genotypes

The most important metric of performance of a genotyping platform is replicability. Independent libraries of a given genotype should cluster in an MDS plot, which is overwhelmingly true across genotypic replicates, but interestingly also across PCR extension time plates (Figures 4, S1, S2). When all four points are not overlapping, PCR extension clusters more tightly than replicate, suggesting some variation in digestion.

**Figure 4.**
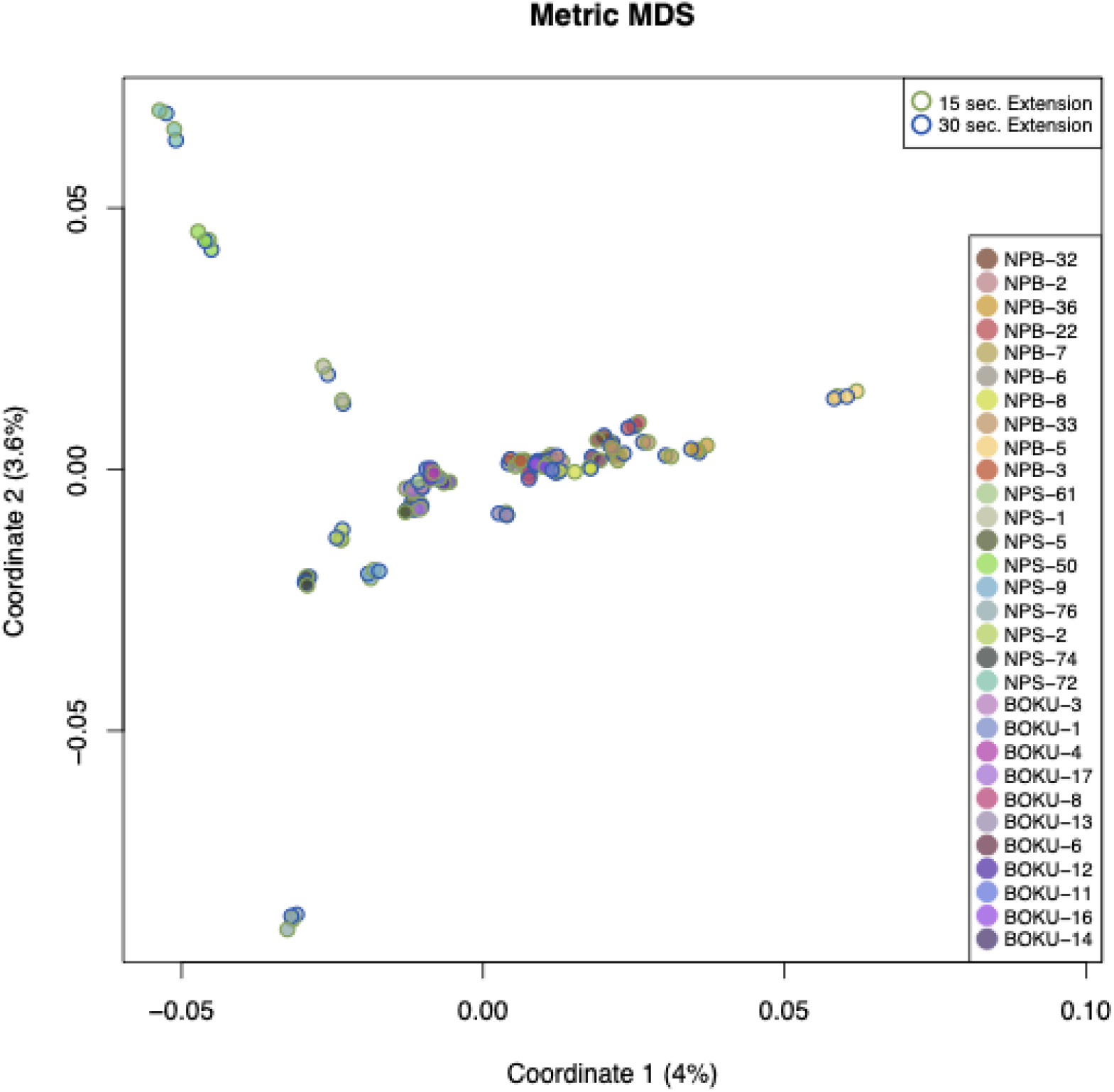
Genotypes cluster across replicates and PCR amplification conditions. Genetic distances calculated from 1.6M SNPs from the dataset “All” mapping 120-mers against the full genome. A genetically diverse, high depth individual from NPS was excluded, because it was driving variation on coordinate 1, decreasing contrast for other samples (Figure S1). The 64-mer version of the dataset performs similarly (Figure S2).

This approach is also excellent at discriminating between genotypes (Figure 5). NPS and NPB, the natural populations from the Bohemian Alps and Alps, respectively, are clearly discriminated on coordinate one, explaining 4.7% of the total variance. Diversity within NPS is greater than between NPB, as defined by increased spread on coordinate 2, explaining 4.2% of the total variance. Interestingly, the managed plot from Austria, BOKU, is central between the two and partially clusters with NPS and partly with NPB.

**Figure 5.**
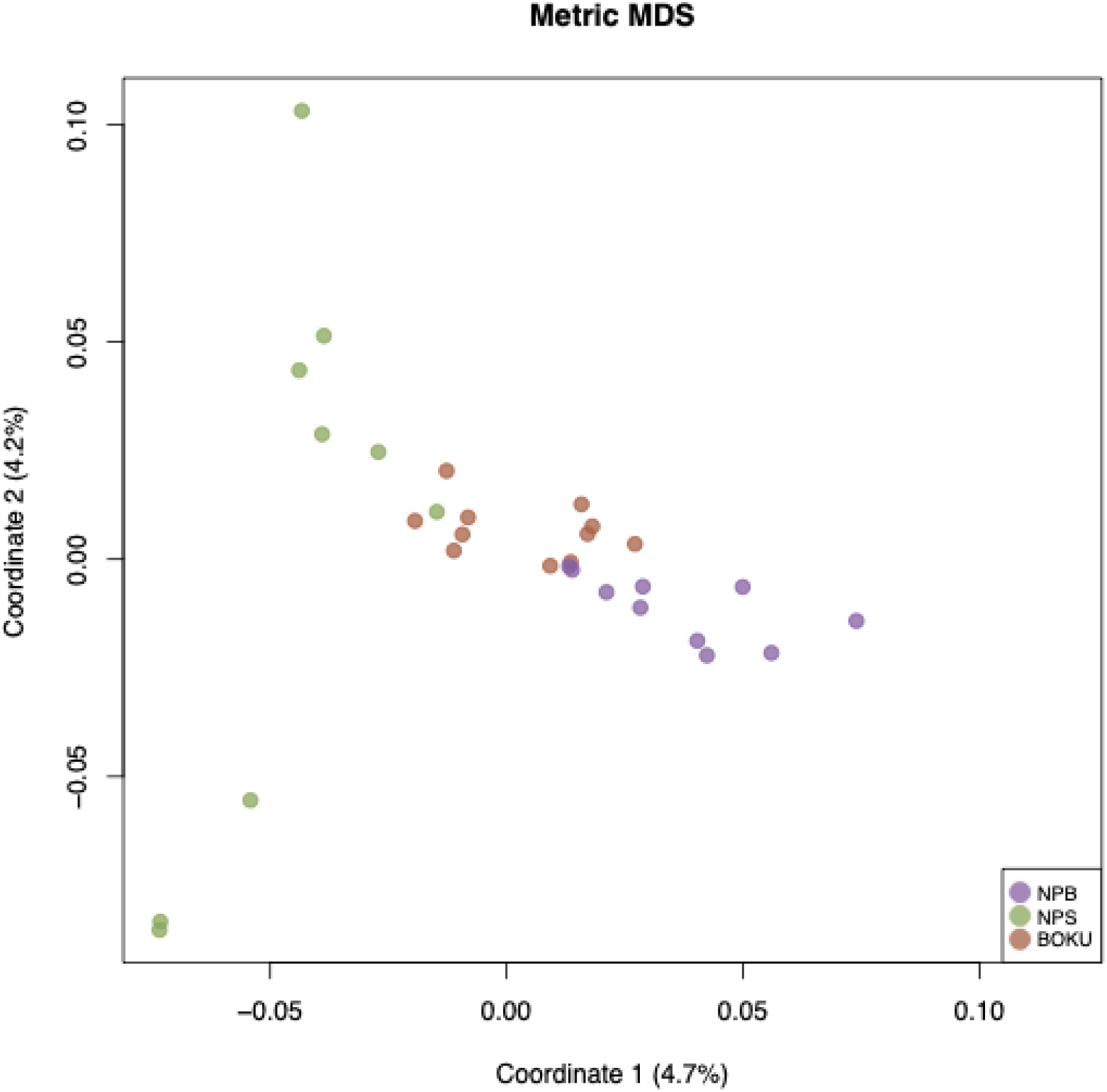
Genotypes cluster by population. ‘BOKU’ is a managed forest whereas ‘NPS’ and ‘NPB’ are natural populations. Genetic distances calculated from 1M SNPs from the dataset “Merge15s” mapping 120-mers against the full genome. A genetically diverse, high depth individual from NPS was excluded, because it was driving variation on coordinate 1, decreasing contrast for other samples.

## Discussion

The greatest advantage of using the above protocol is the ability to massively multiplex samples to generate replicable genotypes at low cost. An Illumina S4 generates 9 billion reads, enough to multiplex 20 plates with 450M reads, or 1920 samples with greater than 5X coverage, for less than 10 euro a sample. Building on the existing GBS framework (Elshire et al., 2011; *Tassel5GBSv2Pipeline*, 2021) has many practical benefits, such as a well documented and tested bioinformatic pipeline. The current pipeline does not account for reverse reads, but this could be added later to the pipeline or with pre-processing.

Switching from *Taq* polymerase to a high-fidelity Phusion allowed us more of the accessible (as indicated by digestion with a methyl-sensitive enzyme) genome but the specifics of the extension length should be tuned for each species. The 15 second extension is expected to amplify 1000bp, which, in spruce, is just inside of the dominant large fragment distribution which we hypothesized to be mostly repetitive, and is supported by the enrichment of CDS in the resulting library. With over a million unbiased SNPs called across only 32 samples, this protocol is well-suited for population genomic analysis and, if the variants are intended for association studies, coverage in functional regions of the genome is comparable to the recent 50K genotyping array (Bernhardsson et al., 2020). As library preparation cost are marginal -- with institutional pricing, we are able to generate libraries for about one euro per sample -- this analysis could be used in conjunction with a genotyping array of choice to ensure back compatibility.

## Supporting information

Supplementary Material 1. GBS. Oligos

Supplementary Material 2. FullProtocol

Supplementary Material 3. ConcatenateReferenceContigs.pl

## Acknowledgements

The authors would like to thank Jaroslav Červenka at NPS, Rupert Seidl and Sebastian Seibold at NPB and Martin Schebeck and Josef Gasch at BOKU for permitting for sample collection. We would also like to thank Asha Jain at the Institute for Genomic Diversity at Cornell for clarification of the standard GBS protocol.

## Author Contributions

MVE and KS conceived of the approach and MVE designed primers. KS, MVE and SS wrote the paper. KS and MVE designed the testing approach and SS carried out extensive early testing of the protocol. VA tested PCR conditions, KS developed robotic 96-plate protocols and KS and LS generated testing libraries. JR extracted DNA and AA was responsible for tissue collection for testing.

## Supplementary material

Supplementary Material 1. GBS. Oligos (.xlsx)

Supplementary Material 2. FullProtocol (.pdf)

Supplementary Material 3. ConcatenateReferenceContigs.pl

**Figure S1:**
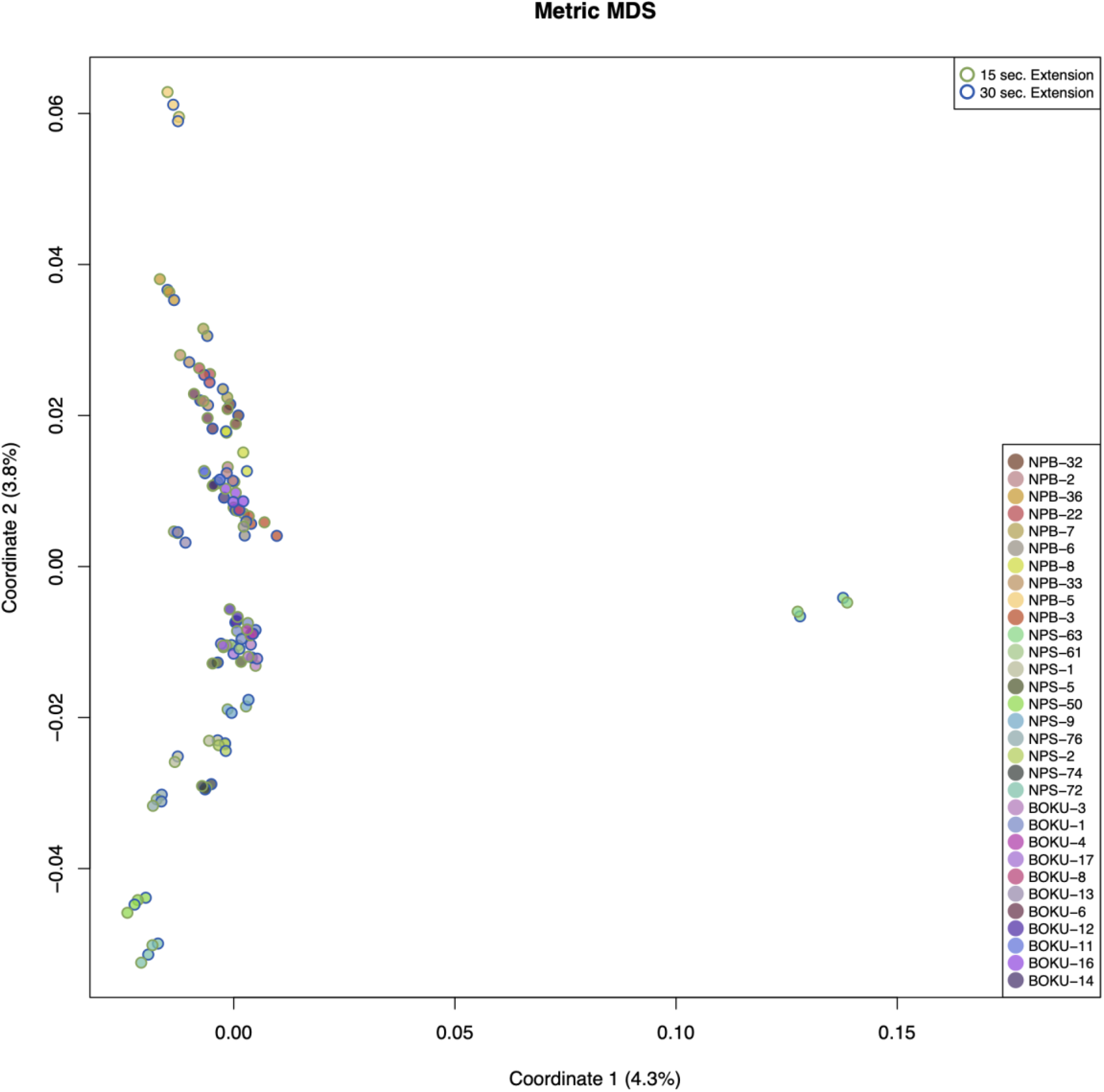
Genotypes cluster across replicates and PCR amplification conditions. Genetic distances calculated from 1.6M SNPs from the dataset “All” mapping 120-mers against the full genome. Including the genetically diverse sample from NPS

**Figure S2:**
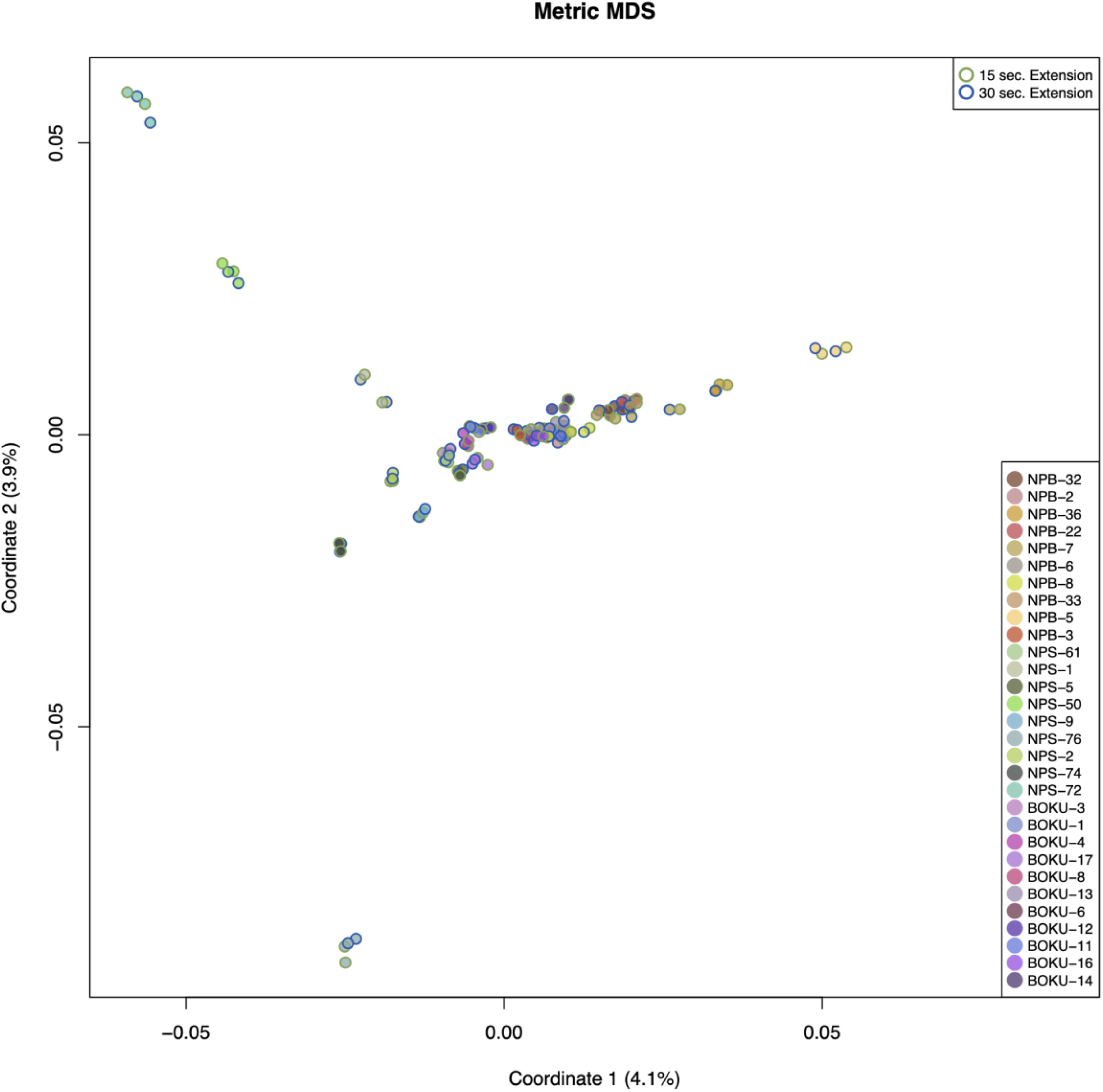
Genotypes cluster across replicates and PCR amplification conditions. Genetic distances calculated from 1.6M SNPs from the dataset “All” mapping 64-mers against the full genome. Including the genetically diverse sample from NPS

