## Supplementary Material 2. FullProtocol for "Multi-GBS: A massively multiplexed GBS-based protocol optimized for large, repetitive conifer genomes"

### PlateGBSApeKI_Published

| | |
|---|---|
| **Category** | Experimental Procedures |
| **Author** | Miguel Vallebueno-Estrada |
| **Version** | Published |

#### Multi-GBS

##### Overview and reagents needed

Protocol used for library building based on restriction of genomic DNA and ligation of adapters over the sticky ends of the restriction sites. As ApeKI is a Methyl sensitive enzyme, it will cut more often over transcriptional active regions and so will therefore be represented as shorter fragments than repetitive regions. During the PCR reaction, longer fragments linearly amplify while short fragments are exponentially amplified acting as a size select.

Reagents needed:
ApeKI (NEB R0643L) + NEB Buffer 3
T4 DNA ligase (NEB M0202L) + ligase buffer
Phusion high fidelity DNA polymerase + buffer (M0530L)
Primer mix GBS_S_UDI## p5+p7
AMPpure (or similar) magnetic beads

Modifications:

- This protocol is based on Asha (personal communication, Cornell Genomic Diversity Facility), Elshire et al 2011 and Wallace et al 2017
- As in the original protocol, we use the methyl sensitive endonuclease ApeKI (NEB R0643L) in order to minimize the repetitive fraction sampled, while still targeting a large fraction of the accessible (functional) regions of the genome. To adjust for this, starting DNA was increased from 100 ng to 150 ng.
- Because we found restriction digestion to be sometimes incomplete at 2 hours, we increased the digestion to overnight (14 hours). NEB buffer 3.1 at the time of testing was unstable. Always check that the digested product follows the expected fragment distribution profile.
- Ligation was increased to 1.5 hours to ensure complete ligation of fragments
- Because the fragment distribution of spruce was shifted to larger digested fragments relative to maize, we use a Phusion high-fidelity polymerase rather than the Taq that is used in the traditional protocol. This necessitates changing PCR conditions. We test here a 15 and 30 second extension time, leading to an expected exponential increase of fragments up to 1000 and 2000 bp respectively. Cycle number was reduced from 18 to 13 to retain library complexity.
- Magnetic bead cleanups were used instead of column cleanups to enable plate-based robotics.
- The volume of the restriction digest and ligation reaction is decreased by half to save costs. Total reagent costs/sample are less than one euro.
- The common adapters were changed to be compatible with TruSeq Illumina Technology (Primer mix GBS_S_UDI## p5+p7), allowing for plate-level multiplexing.

#### Samples

1. Make GBS plate with 3.6 ng of GBS-adapter pairs (common + specific) and dry down.
2. Add **150 ng** of the sample genomic DNA and dry in SpeedVac

#### Restriction

**Restriction Master Mix**

| Reagent | Stock concentration (M) | Final concentration (M) | Volume per sample (ul) | one plate/98 samples (ul) |
|---|---|---|---|---|
| NEB Buffer 3.1 (LOT: XXXXXXXXXXX) | 10X | 1X | 1 | 98 |
| ApeKI (NEB R0643L) (LOT: XXXXXXXXXXX) | 4U/ul | 1U/100ng DNA | 0.125 | 12.25 |
| $H_2O$ | | | 8.875 | 869.75 |
| **Final Volume** | | | 10 | 1960 |

1. Prepare the Restriction master mix.
2. Add **10 µl** of digestion master mix to each sample tube.
3. Spin down and incubate reaction at **75°C** for **14 hours** then hold at 4°C until ready to ligate. Lid has to be pre-heated, PCR plate has to be stabilized with a silicone compression mat (Axymat).

#### Ligation

**Ligation master mix**

| Reagent | Stock concentration (M) | Final concentration (M) | Volume per sample (ul) | one plate/100 samples (ul) |
|---|---|---|---|---|
| Ligase Buffer (LOT: XXXXXXXXXXX) | 10X | 1X | 2.5 | 250 |
| T4 DNA ligase (NEB M0202L) (LOT: XXXXXXXXXXX) | 400000U/ml | 400 Cohesive NEB Units | 0.5 | 50 |
| $H_2O$ | | | 12 | 1200 |
| **Final Volume** | | | 15 | 1500 |

Caution: if you do not use T4 ligase from New England Biolabs, activity is usually expressed in Weiss Units (1 Weiss Unit = 67 NEB Cohesive End Ligation Units).

1. Add 15 **µl** of ligation master mix to each sample tube containing the 10 ul from the previous step.

2. Spin down and ligate at **22°C** for **1.5 hour**. Kill the ligase by heating to **65°C** for **20 min** then hold at 4C. PCR plate has to be stabilized with a silicone compression mat (Axymat).

#### Clean up beads 2.0X

Use robot protocol OR if need to do by hand:

1. Add **100 µl** of beads (2.0X volumes) to the PCR tube and mix
2. Mix thoroughly by pipetting
3. Incubate 5 min at room temperature
4. Place the tubes into a magnetic rack and wait 2 min.
5. Without disturbing the pellet discard supernatant
6. Wash with **200 ul** of **80% EtOH**
7. Without disturbing the pellet discard EtOH
8. Wash with **200 ul** of **80% EtOH**
9. Without disturbing the pellet discard EtOH
10. Dry sample for 2 min.
11. Elute in **50** µl of **EB** Elution buffer.
12. Incubate 5 min
13. Place the plate into a magnetic rack and wait 2 min.
14. Transfer the supernatant into a new PCR plate.
15. Label plate with library number and add **"Ligation"**legend to it.
16. store at -20 °C if needed.

#### PCR to add TruSeq adapters

PCR Rxn

| Reagent | Stock concentration | Final concentration | Volume per sample(ul) | one plate/100 samples (ul) |
|---|---|---|---|---|
| Sample | | | 13 | - |
| 5X Phusion buffer | 5X | 1X | 10 | 1000 |
| p5 adapter **UDI_XX** | 100 µM | 0.5 µM | 0.125 | 12.5 |
| p7 adapter **UDI_XX** | 100 µM | 0.5 µM | 0.125 | 12.5 |
| dNTPs (100mM) | 25mM /each | 0.2 mM | 0.4 | 40 |
| H20 | | | 25,35 | 2525 |

| Reagent | Stock concentration | Final concentration | Volume per sample(ul) | one plate/100 samples (ul) |
|---|---|---|---|---|
| Phusion polymerase (LOT: XXX) | unknown | 1U | 1 | 100 |
| Final Volume | | | 50 | 1250 |

**Keep on ice! Phusion is not a hotstart!**

1. From master mix, do not put in multichannel reservoir but rather split into a pcr strip tube because very viscous (250/tube for 100 samples)
2. Put Elution plate of DNA onto alpaqua ring magnet and let sit 3 minutes
3. Transfer 13ul of ligated, digested product to an unskirted PCR plate in holder and add 37ul of PCR master mix (50ul rxns)
4. Add PCR plastic sticky top, mix, spin and move to thermocycler
5. Amplify on a thermocycler using the following program :

98 ◦C for 30 sec
98 ◦C for 10 sec
**13X** 65◦C for 15 sec
72 ◦C for 15 sec
72 ◦C for 5 min
4 ◦C hold

Save remaining ligation product in freezer

#### Clean up beads 1.1X

With robot OR if need to do by hand:

1. Add **55 µl** of beads (1.1X volumes) to the PCR tube and mix ***This is still fairly permissive. Can size select harder at this step.
2. Mix thoroughly by pipetting
3. Incubate 5 min at room temperature
4. Place the tubes into a magnetic rack and wait 2 min.
5. Without disturbing the pellet discard supernatant
6. Wash with **200 ul** of **80% EtOH**
7. Without disturbing the pellet discard EtOH
8. Wash with **200 ul** of **80% EtOH**
9. Without disturbing the pellet discard EtOH
10. Dry sample for 2 min.
11. Elute in **50** µl of **EB** Elution buffer.
12. Incubate 5 min
13. Place the plate into a magnetic rack and wait 2 min.
14. Transfer the supernatant into a new PCR plate.

15. Label tube with library number and add **"GBS Lib"** legend to it.

Store at -20 °C (this is the final build library)!

#### Expected fragment size distribution

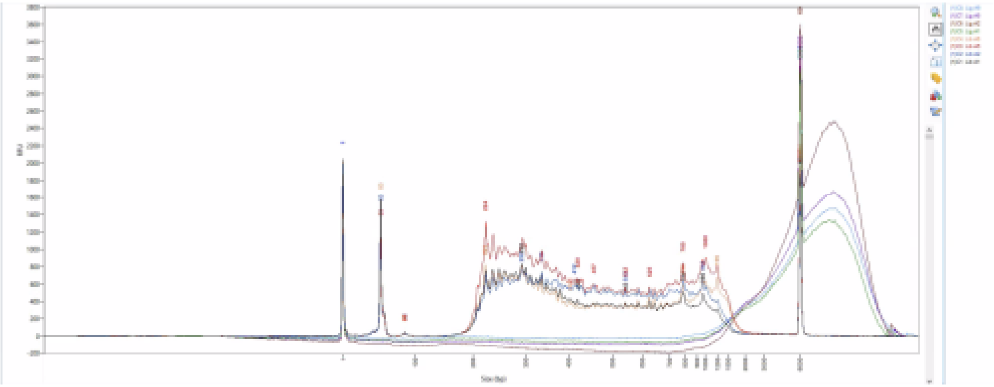

Screen Shot 2021-08-24 at 09.55.33.png

#### Attachments

| FileID | Name |
|---|---|
| 4531 | illumina-adapter-sequences-1000000002694-11.pdf |
| 4533 | Elshire et al 2011 GBS.PDF |

*This procedure was originally created by* ***Miguel Vallebueno***

Powered by eLabNext
